# Nanoscale Mapping Reveals Periodic Organization of Neutrophil Extracellular Trap Proteins

**DOI:** 10.1101/2025.07.28.665103

**Authors:** Moritz Winkler, Jan Schmoranzer, Garth Lawrence Burn, Niclas Gimber

**Author notes:** Correspondence should be addressed to Niclas Gimber and Garth Burn.

## Abstract

Neutrophils are essential cells of the innate immune system that release Neutrophil Extracellular Traps (NETs) via NETosis. This specialized cell death pathway results in the extrusion of decondensed chromatin into the extracellular space. NET formation participates in antimicrobial defense and coagulation but dysregulated NETosis can contribute to neoplasticity, coagulopathy, and autoimmune disease. Here, we present a workflow combining optimized sample preparation, super-resolution imaging and quantitative bioimage analysis to investigate the nanoscale organization of NETs. Our newly developed analysis tool, NanoNET, facilitated the automated identification of DNA filaments and the analyses of NET binding proteins along those filaments through auto- and cross-correlation. We identify specific NET proteins, including neutrophil elastase (NE) and proteinase 3 (PR3), that are bound along DNA strands in a periodic fashion that mirrored nucleosomes. Cross-correlation of NE and PR3 with nucleosomes revealed that these proteins were highly colocalized suggesting that they dock onto or around nucleosomes. The presented workflow and analysis tools represent a significant methodological advance for studying protein distributions along NETs and along filamentous structures in general. Understanding how NET filaments are organized is a critical step toward elucidating both their formation and function.

## INTRODUCTION

Neutrophils play a pivotal role in innate immunity during septic and aseptic injury ^1,2^. One key effector mechanism of neutrophils is the formation of Neutrophil Extracellular Traps (NETs) via a cell death program called NETosis ^3^. During NETosis, nuclear chromatin is decondensed and transformed into cell-free scaffolds by cytoplasmic effector proteins. These atypical chromatin structures contain modified nucleosomes and are decorated with a distinct set of cytoplasmic proteins ^4,5^. Both exogenous and endogenous stimuli can lead to NETosis ^6^. Once initiated, NET formation proceeds via several discrete yet overlapping processes which includes the translocation of cytoplasmic proteins into the nucleus, nuclear envelope breakdown, the intermixing of decondensed chromatin with the cytoplasm and finally, the rupturing of the plasma membrane which releases NETs into the extracellular space ^7^. When NETs are released appropriately, they play an important role in maintaining homeostasis by trapping and killing microbes or initiating coagulation, however, the unbridled release of NETs is associated with neoplasticity, coagulopathy and autoimmune disease ^8^. Despite the growing clinical relevance of NETs, the underlying nanoscale organization of NET proteins along extruded DNA filaments remains unknown. NET filaments stained with both DNA dyes and antibodies appear in a contiguously labeled manner when employing conventional diffraction limited microscopy methods that cannot resolve crowded overlapping fluorophores ^9,10^. To overcome this limitation, we used three different super-resolution techniques ^11^ in combination with our newly designed quantitative software toolbox that we called NanoNET.

Using NanoNET, we were able to identify DNA strands and subsequently investigate periodic binding patterns of different NET associated proteins in an automated way. By offering detailed protocols and software, we aim to make this workflow scalable for studying the localization of proteins along NET DNA filaments. Understanding the nanoscale organization of proteins on NETs is a key step toward unraveling the structure-function relationships that underpin NET biology.

## RESULTS AND DISCUSSION

### Imaging and analysis workflow for Neutrophil Extracellular Traps

To enable reliable imaging of individual NET filaments without excessive DNA aggregation or disruption of the nano-structure, we have developed a protocol that safeguards DNA filaments from extensive aggregation and makes them compatible with super-resolution imaging (Figure 1a). After seeding freshly isolated neutrophils onto coverslips, NET formation was induced using the mitogen Phorbol 12-Myristate 13-Acetate (PMA), followed by a specific fixation protocol and mounting procedure. A detailed description of the protocol is provided in the methods section. We used the DNA intercalating dye YOYO-1 to stain the NET backbone, due to its high quantum yield and good signal-to-noise ratio YOYO-1 was ideal for NET detection (Figure S1). To automate the analysis of thousands of uniformly sized NET fragments, we developed the NanoNET Toolbox (https://github.com/ngimber/NanoNET), which efficiently segments DNA strands (Figure 1b, d) and produces auto- and cross-correlation plots for NET-associated proteins (Figure 1c, d). Using antibodies directed against NET specific proteins, we were able to analyze the periodicity and colocalization of those proteins along the DNA-labeled NET filaments. Utilizing three different super-resolution microscopy techniques, and by generating auto- and cross-correlation histograms, we demonstrate here that a subset of highly abundant ^4^ NET binding proteins are periodically distributed and colocalize with nucleosomes.

**Figure 1:**
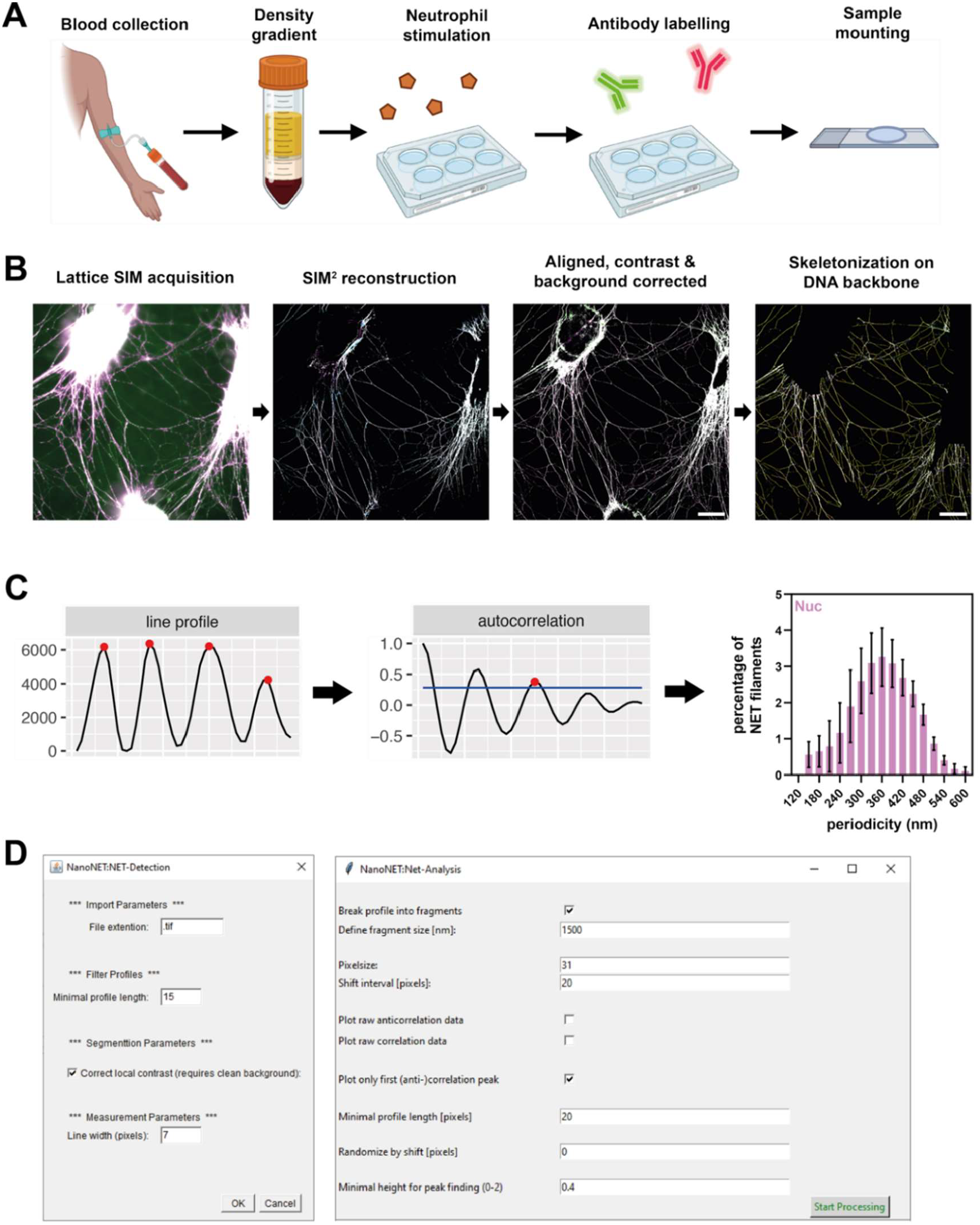
Nanoscale protein mapping of NET samples. **(a)** Schematic of NET preparation for super-resolution microscopy. Neutrophils were purified by a two-step density gradient from fresh blood samples collected from healthy donors. NET formation was stimulated with PMA on glass coverslips followed by fixation, antibody labelling and sample mounting (details are provided in the methods). **(b)** Imaging and processing pipeline. SIM images were acquired and reconstructed using the Zeiss SIM^²^ module. Multicolor channels were aligned and images were background corrected. Regions with single thin filaments were automatically processed using ‘NanoNET-Detection’ for the detection of single NET filaments based on DNA signal. Scale bar = 10µm. **(c)** ‘NanoNET-Analysis’ was used to extract intensity line profiles along skeletonized NET filaments as equally sized fragments, to perform the autocorrelation analysis of NET protein signals and to generate periodicity histograms. **(d)** NanoNET toolbox for automated NET-Detection (left) and NET-Analysis (right).

### SIM Reveals Periodic Distribution of Nucleosomes on NETs and Distinct Binding Modes of Neutrophil Serine Proteases

We used multi-channel structured illumination microscopy (SIM) to investigate the nanoscale organization of NET proteins and nucleosomes (as potential interaction hubs) along NET filaments. SIM provides a 2-fold gain of resolution over conventional (confocal) imaging, both in the lateral (x,y) and axial (z) dimension, and allows the use of conventional fluorophores ^12^. Additionally, SIM does not require very high laser powers and offers shorter acquisition times compared to other super-resolution techniques (e.g. SMLM and STED), making this technique perfectly suited to initially screen several NET associated proteins on the nano-scale, while maintaining a medium-to-high throughput ^11^. We first focused on the Neutrophil Serine Proteases (NSP) NE and CATG. NSPs are a set of highly conserved immune cell restricted proteases that are expressed mainly in neutrophils and have been shown to be important immune effectors by cleaving bacterial virulence factors, remodeling extracellular matrix, regulating pro-inflammatory mediators and, interestingly, are also implicated in NET formation ^13,14^. Both NE and CATG have previously been reported to bind to NETs ^4^. NETs were co-labeled with YOYO-1 (DNA-backbone), the nucleosome antibody PL2.3 (recognizes a conformational H2A-H2B-DNA epitope) and antibodies against NE and CATG. By applying SIM we found that nucleosomes, NE and CATG localize as globular puncta along individual NET filaments (Figure 2a, Figure S1), reminiscent of the globular structures seen on NETs by scanning electron microscopy (SEM) ^15^. To comprehensively analyze the spatial distribution of NET proteins along the DNA-backbone, including tests for potential periodic distributions, we designed the toolbox NanoNET. Using NanoNET allowed us to generate spatial autocorrelograms from a high number of NET protein signals (e.g. nucleosomes, NE and CATG) along the automatically detected DNA-backbone. For each protein, we determined the lag distance of the first peak within the autocorrelogram to generate a periodicity histogram of a large number of individual NET protein profiles (see Methods; Figure 2b, c). Using Gaussian fitting, we extracted the predominant periodicity from the histogram. We found that NE and nucleosomes exhibited similar periodicities of 370 nm and 383 nm, respectively (Figure 2d), reflecting comparable spacing between individual nano-domains of both proteins along the filamentous meshwork of NETs. Similarly, proteinase 3 (PR3) another NSP family member displayed a strong periodicity at 375 nm (Figure S2). In contrast, CATG displayed a weak periodicity (low peak), albeit with a similar spacing as nucleosomes, NE and PR3. Altogether, this might indicate that NE and PR3 depend on a nearby nucleosome to bind to NETs, while CATG follows a different binding scheme.

**Figure 2:**
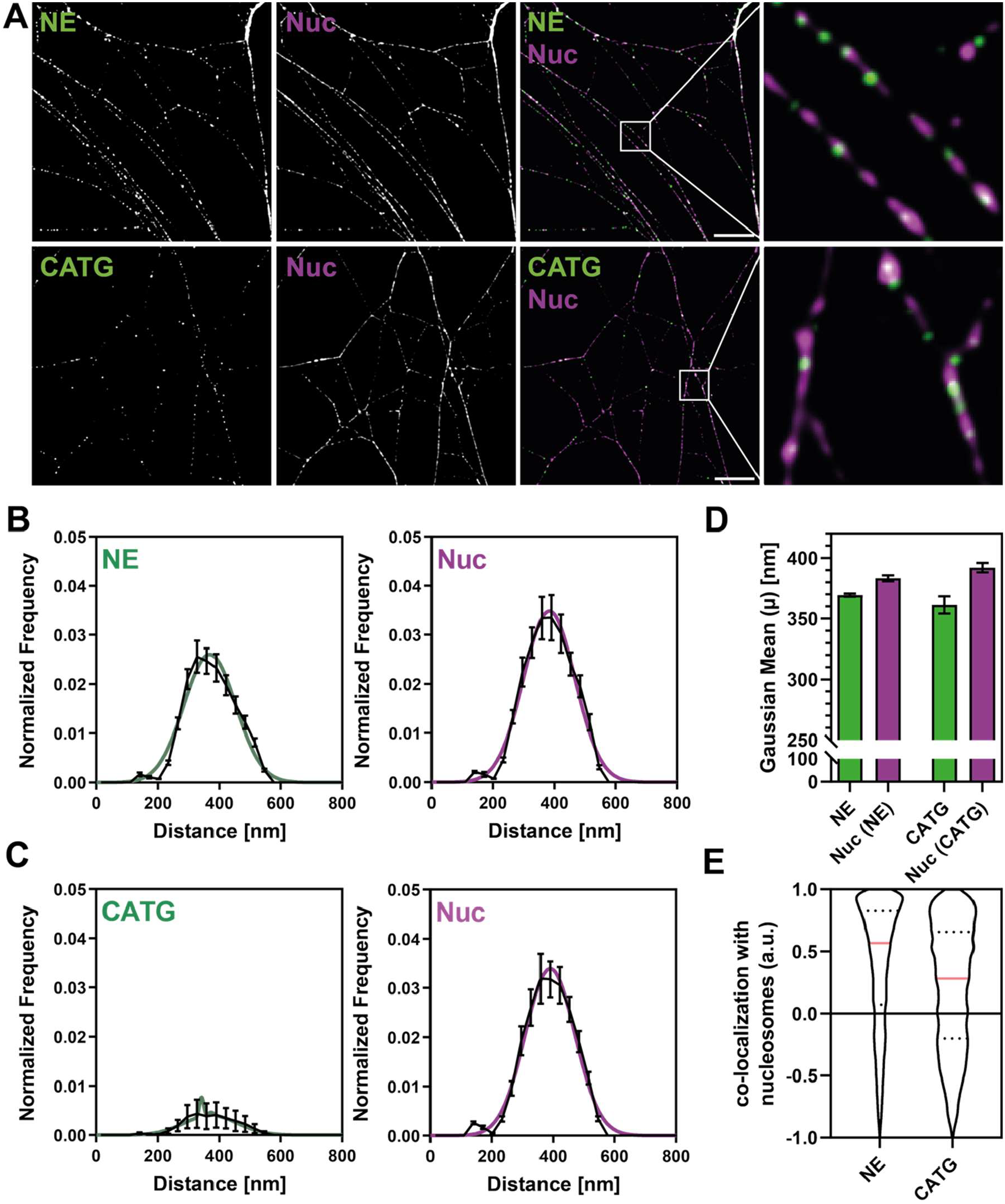
SIM reveals periodic distribution of nucleosomes on NETs and distinct binding modes of different NSPs. Co-labeling of NSPs with nucleosomes (Nuc, mAB PL2.3) on NETs. Data from five independent donors. Analyzed line profiles (1.5 μm each): NE = 60,420; CATG = 66,205. **(a)** Representative SIM images of NE (top) and CATG (bottom) co-labeled with nucleosomes. Scale bar: 3 μm; inset box: 2 × 2 μm. **(b, c)** Periodicity histograms for NE, CATG, and Nuc along NETs. Averages of Gaussian fits are shown as colored lines. Note the pronounced periodicity of NE and Nuc (prominent peaks), in contrast to CATG, which lacks prominent periodicity. Mean ± SEM shown as black lines. **(d)** Centers of the Gaussian fits in (b, c), reflecting the predominant periodicity of each protein. Mean ± SEM. **(e)** Cross-correlation analysis of NE or CATG with nucleosomes, replotted from Figure S6. NE shows strong colocalization with nucleosomes, whereas CATG displays a multimodal distribution. Analysis only includes double-labeled NET fragments (NE: 32.6%; CATG: 11.2%). Median ± quartiles.

To test whether all three NSPs co-localize with nucleosomes, we analyzed their cross-correlation with nucleosomes using NanoNET. Our analysis revealed that NE and PR3 were highly cross-correlated / colocalized with nucleosomes (Figure 2e, Figure S6), while CATG exhibited a lower cross-correlation with nucleosomes compared to NE and PR3 (Figure 2e, Figure S6). These data corroborate our previous findings when comparing the periodicity of NE and PR3 with nucleosomes suggesting that these two proteins directly or indirectly associate with or around nucleosomes, while CATG follows a different binding mode. We validated the periodicity analysis by comparing either two NE antibodies (monoclonal vs polyclonal) or two nucleosome markers (PL2.3 vs 3D9, an antibody against the cleavage site of NET-associated nucleosomes on the H3 tail ^16^; Figure S3). In both conditions, two markers against the same target revealed very similar periodicity in terms of spacing and frequency, demonstrating the robustness of our assay and analysis pipeline.

Additionally, we applied our spatial analysis to other NET proteins that have been recently identified by mass spectrometry. Human neutrophil peptide 1 (HNP-1), catalase (CAT), calgranulin B (S100A9), and transaldolase 1 (TALDO1) displayed no clear periodicity and, except for HNP-1, exhibited no significant colocalization with nucleosomes (Figures S4–S6).

Altogether, our results suggest that periodicity is a distinct feature of specific NSPs, particularly NE and PR3, and is closely associated with their colocalization with nucleosomes. Other NET-associated proteins display less periodic signal or do not colocalize readily with nucleosomes. This differential organization may reflect distinct ways in which different proteins interact with NETs.

### STED Microscopy Confirms Periodic Binding Patterns of NET Proteins

The SIM system we used provides a lateral resolution of ∼100 nm, whereas the available STED setup can resolve structures as small as 50–75 nm. Therefore, we used STED (Figure S7) to validate the periodicity observed with SIM and to search for potentially smaller underlying structures without the need for image reconstruction ^17^. STED images of nucleosomes co-labeled with NE or PR3 revealed a highly periodic distribution of both proteins along NET filaments, consistent with the patterns observed using SIM, while CATG showed no obvious periodicity (Figure 3a, Figure S8a). We analyzed the images using NanoNET and obtained periodicity histograms that revealed pronounced peaks for NE, PR3 and nucleosomes, confirming their periodic organization (Figure 3b-d, Figure S8b,c). CATG, however, lacked any periodicity in STED which might be due to the lower sensitivity (stronger photobleaching) of STED compared to SIM (Figure 3c). Both super-resolution methods revealed highly periodic distributions for NE, PR3 and nucleosomes, but a predominantly aperiodic distribution of CATG.

**Figure 3:**
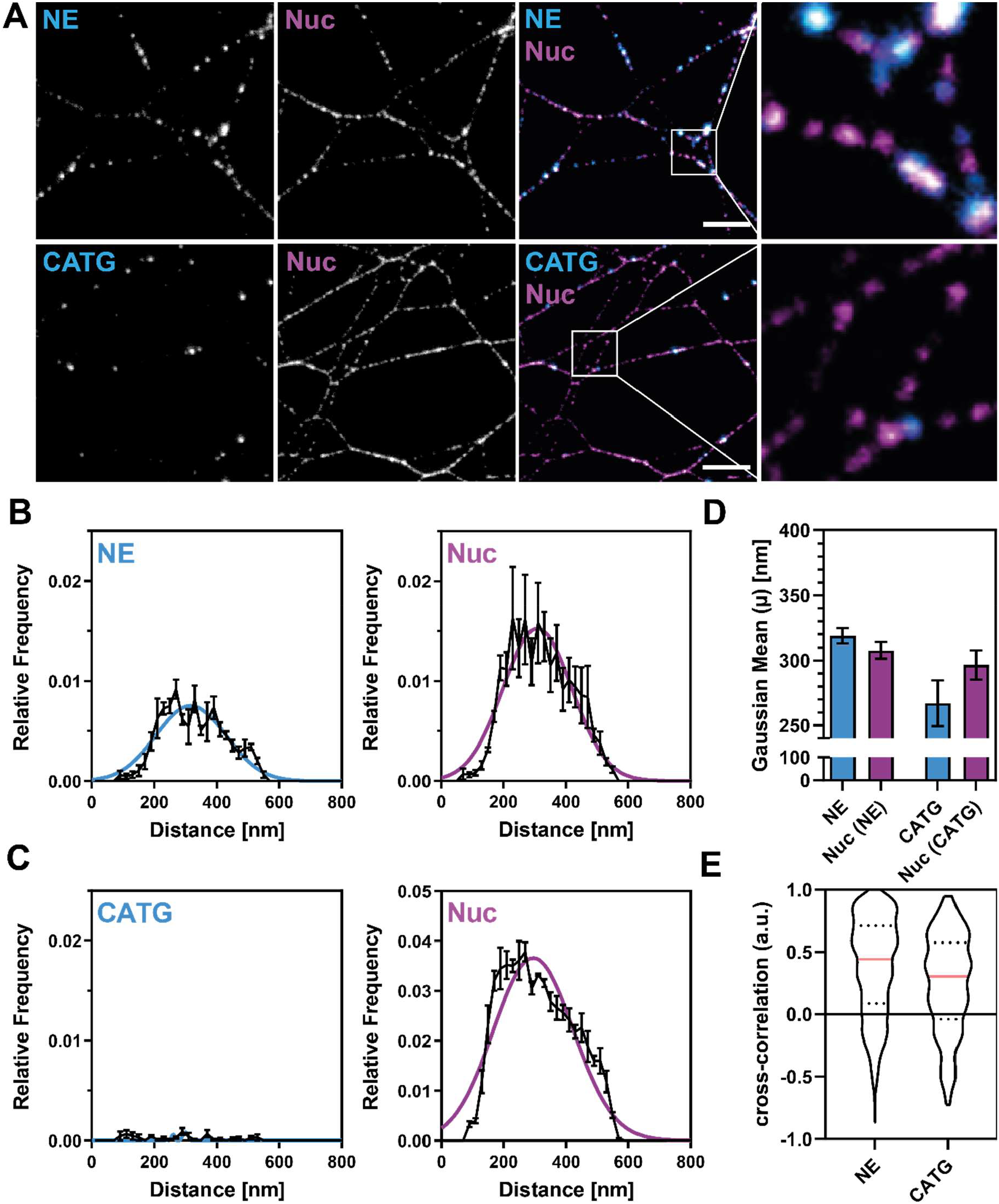
STED microscopy validates the periodic binding patterns of proteins on NETs and the colocalization of NE and Nuc. Co-labeling of neutrophil serine proteases with nucleosomes (Nuc mAB PL2.3) on NETs. Data from three independent donors. Analyzed line profiles (1.5 μm each): NE: = 10,974; CATG = 9,219. **(a)** Representative STED images of NE (top) and CATG (bottom) co-labeled with nucleosomes. Scale bar: 2 μm; inset box: 2 × 2 μm. **(b, c)** Periodicity histograms for NE, CATG, and Nuc along NETs validate the pronounced periodicity of NE and Nuc (Figure 2). Averages of Gaussian fits are shown as colored lines. Mean ± SEM shown in black lines. **(d)** Centers of the Gaussian fits in (b, c), reflecting the predominant periodicity of each protein. Mean ± SEM. **(e)** Cross-correlation analysis of NE or CATG with nucleosomes, replotted from Figure S9. NE shows strong colocalization with nucleosomes, whereas CATG displays a multimodal distribution. Analysis includes only double-labeled NET fragments (NE: 16.2%; CATG: 1.5%). Median ± quartiles.

By using NanoNET for the cross-correlation analysis of STED imaging, we confirmed the strong colocalization of NE and PR3 with nucleosomes observed with SIM, while CATG showed no significant colocalization with nucleosomes (Figure 3e, Figure S9). In summary, STED strengthens the finding that NE and PR3 share periodic binding sites with nucleosomes, while CATG follows a distinct binding mode without significant periodicity. Additionally, our results demonstrate that NanoNET can be used to robustly analyze images at different scales from independent super-resolution methods.

### Single-Molecule Localization Microscopy (SMLM) Reveals Sub-Diffraction Periodicity of NE and Nucleosomes on NETs

The periodicity of NET-associated proteins observed in SIM and STED and the colocalization with nucleosomes suggests that some NET proteins are highly organized along filaments and that this organization, in turn, may be afforded by the presence of a nucleosome. However, due to their resolution limits, both techniques might miss finer spatial (periodic) distributions. To overcome this limitation, we employed direct stochastic optical reconstruction microscopy (dSTORM) ^18^, a single-molecule localization microscopy method that achieves a resolution of ∼ 20 - 30 nm on our SMLM system^19^.

dSTORM experiments on NE and nucleosomes confirmed that both proteins form periodic clusters along individual NET filaments (Figure 4a). We quantified these periodic structures by generating periodicity histograms with the ‘NanoNET-Analysis’ module on NET samples from multiple donors. In contrast to the single-Gaussian distribution of periodicities observed in SIM and STED, dSTORM revealed a bimodal distributed periodicity of both proteins, indicating two underlying distributions (Figure 4b). A two-component Gaussian fit of the data revealed two predominant periodicities, approximately ∼150 nm and ∼300 nm for both proteins, which represent integer multiples of each other (Figure 4c).

**Figure 4.**
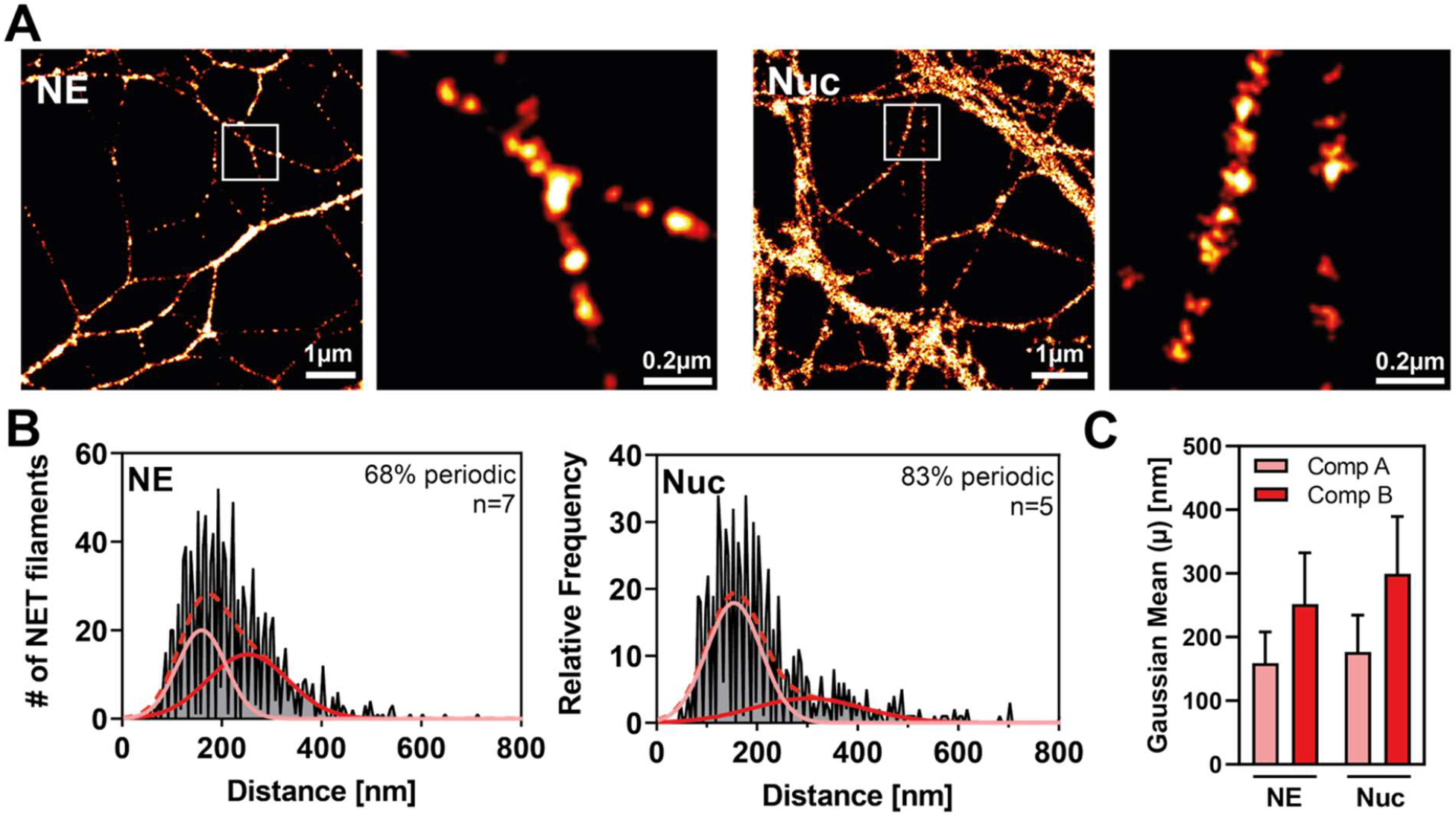
SMLM reveals sub-diffraction periodicity of NE and Nuc on NETs. Co-labeling of NE and nucleosomes (Nuc, mAB PL2.3) on NETs using SMLM (dSTORM). Data from 7 (NE) and 5 (Nuc) independent donors; analyzed NET fragments: NE = 1,084, Nuc = 773 **(a)** Representative images. NE and Nuc form clusters along single NET filaments. Scale bar = 1 µm. **(b)** Periodicity histogram of the proteins displayed in (a). The exceptional resolution of SMLM reveals a bi-modal periodicity (validated by Akaike information criterion), in contrast to the single-Gaussian distribution observed in SIM/STED (Figure 2 and 3). Averages of the two-component Gaussian fits (solid red line) and individual components (dashed red lines) are displayed. **(c)** The two predominant periodicities (centers of both components from (b) are plotted. Means ± SEM.

SIM and STED likely failed to detect the high-frequency component of the bimodal distribution due to their resolution limits, thereby missing periodic clusters with ∼150 nm spacing. The higher resolution of SMLM resolved these missing structural repeats, offering a more detailed view of NET organization.

## CONCLUSION

In this study, we provide an integrated workflow of sample preparation, super-resolution imaging and bioimage analysis to investigate the nanoscale organization of NET associated proteins. While SIM, STED, and SMLM are all powerful super-resolution methods that allow imaging with resolutions beyond the diffraction limit, each method has its own strengths and limitations regarding resolution, acquisition speed and image processing. By combining these three complementary methods and by analyzing the data with our newly developed NanoNET toolbox, we gain insight into the nano-scale organization of NET associated proteins along single NET filaments. Our results demonstrate that a subset of NET binding proteins like NE and PR3 are arranged in a highly periodic fashion along NET filaments, mirroring the periodicity of nucleosomes on NETs. Moreover, both NE and PR3 are highly cross-correlated with nucleosomes, leading us to believe that these proteins likely depend either directly or indirectly on the positioning of nucleosomes on NETs to bind. Interestingly, despite ∼90 % sequence homology to PR3 and NE, the neutrophil serine protease CATG displays a more aperiodic distribution and a weaker correlation with nucleosomes than PR3 and NE. NE, PR3, and CATG are all highly positively charged (pI values of 10.5, 9.4, and 8.9, respectively) and therefore prone to interact with the DNA backbone’s negatively charged phosphate groups. However, CATG binds double-stranded DNA about 20 times more tightly than NE (Kd ≈ 35 nM vs 607 nM) ^20^, showing that net charge alone cannot predict their DNA affinities. NE might be retained on NETs through additional interactions, such as with histones or other nucleosome-associated proteins (Figure S10). CATG on the other hand, might bind directly to inter-nucleosomal DNA given its high affinity for double stranded DNA, consistent with our results. Alternatively, there may also be quaternary DNA structures that influence binding, that we could not account for ^21^.

The average distance between nucleosomes reported within the nucleus is between 10-30 nm ^22,23^. In contrast, our results show that in NETs this distance increases to ∼150 nm, suggesting that nucleosomes may be disassembled and evicted during NETosis. This apparent reduction in the total number of nucleosomes on NETs would be consistent with ATAC-see and ATAC-seq data acquired from resting neutrophil chromatin versus NETs showing largescale changes in nucleosome occupancy and configuration on chromatin as NETosis proceeds ^24^. The eviction of nucleosomes from NETs might be functionally significant allowing for decondensation of DNA as well as providing docking sites for proteins that decorate NETs independently of nucleosomes but dependently on DNA.

In summary, our nano-scale analyses show that some NET-associated proteins are specifically organized along individual NET filaments. Our newly designed open access tools and protocols will facilitate future research on how proteins organize along NET filaments. This nano-organization of proteins within NETs will help generate new hypotheses with respect to NET formation and function. In addition to NETs numerous other filamentous structures, including cytoskeletal elements or neuronal structures, could be analyzed for periodic features using NanoNET. We regard NanoNET as a general tool that could be applied to study distributions of fluorescent markers on any filamentous structure at any scale.

## ASSOCIATED CONTENT

### Supporting Text

Detailed experimental methods and material.

### Supporting Figures

Supporting Figure S1: Triple-color SIM of NET proteins; Supporting Figure S2: SIM of PR3 and nucleosomes; Supporting Figure S3: Dual-color controls (SIM); Supporting Figure S4: SIM of granular and cytosolic proteins; Supporting Figure S5: Periodicity analysis of granular and cytosolic proteins (SIM); Supporting Figure S6: Colocalization analysis of NET proteins with nucleosomes (SIM); Supporting Fig 7: Multi-color STED microscopy of NETs; Supporting Figure S8: Multi-color STED microscopy of PR3 and nucleosomes; Supporting Figure S9: Colocalization analysis of NET proteins with nucleosomes (STED); Supporting Figure S10: Schematic binding model of different neutrophil serine proteases.

### Supplementary Tables

Supplementary Table 1: Primary antibodies; Supplementary Table 2: Secondary antibodies; Supplementary Table 3: Dyes & kits;

## AUTHOR INFORMATION

**Corresponding Authors:** * Phone: 0049 30 450 536333 and

### Author Contributions

G.B. conceived the project. G.B. and N.G. supervised the project. G.B., M.W., N.G. and J.S. designed the project and conceptualized the data. M.W. and N.G. performed experiments and analyzed data. N.G. designed and implemented the software tool NanoNET. N.G., G.B. and M.W wrote the manuscript. J.S. gave advice on the structure and content of the manuscript.

### Notes

The authors declare no competing financial interest.

## ACKNOWLEDGEMENT

We thank the AMBIO imaging facility and staff for their help with training and support throughout the project. We thank Olivia Majer for SIM training and access to SIM microscopy. We thank Jörg Piontek and Ayk Waldow for their help, input and access to STED microscopy, and we would especially like to thank Jan Göing for important discussions concerning analysis. We gratefully thank Arturo Zychlinsky for providing funding and input for the project.

## ABBREVIATIONS

SMLM: single-molecule localization microscopy
STORM: stochastic optical reconstruction microscopy
STED: stimulated emission depletion
SIM: structured illumination microscopy
SEM: scanning electron microscopy
NET: neutrophil extracellular traps
NSP: neutrophil serine proteases
NE: neutrophil elastase
CATG: cathepsin G
PR3: proteinase 3
HNP-1: human neutrophil peptide 1
CAT: catalase
S100A9: calgranulin B
TALDO1: transaldolase
PMA: Phorbol 12-Myristate 13-Acetate
CLSM: Confocal laser scanning microscopy.

