## Supplementary Information for "Nanoscale Mapping Reveals Periodic Organization of Neutrophil Extracellular Trap Proteins"

### SUPPORTING FIGURES

**Supporting Figure S1.** Triple-color SIM of NET proteins.

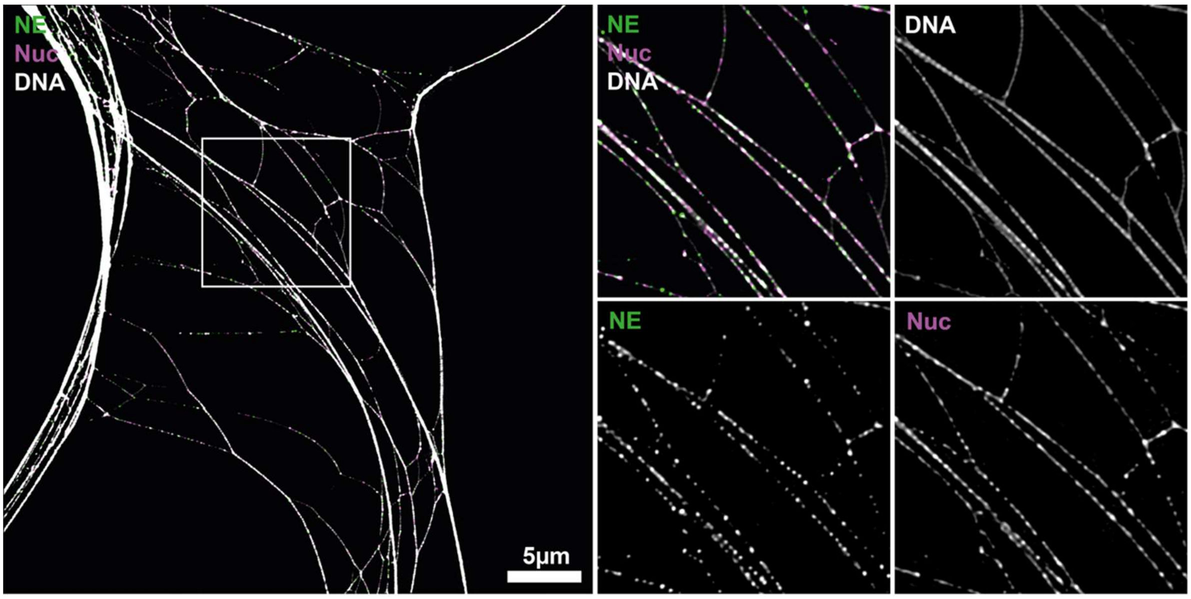


**Supporting Figure S1: Triple-color SIM of NET proteins.** The DNA backbone (white) was stained with the intercalating dye YOYO-1 and co-labeled with antibodies against NE (green) and Nuc (purple). The overview image (left) reveals the characteristic web-like structure of NETs, including individual filaments and unresolved dense regions, which were excluded from analysis. Enlarged regions (right, 10 × 10 µm) highlight the densely labeled DNA backbone, while NE and Nuc appear in periodic clusters along the filaments.

**Supporting Figure S2.** SIM of PR3 and nucleosomes.

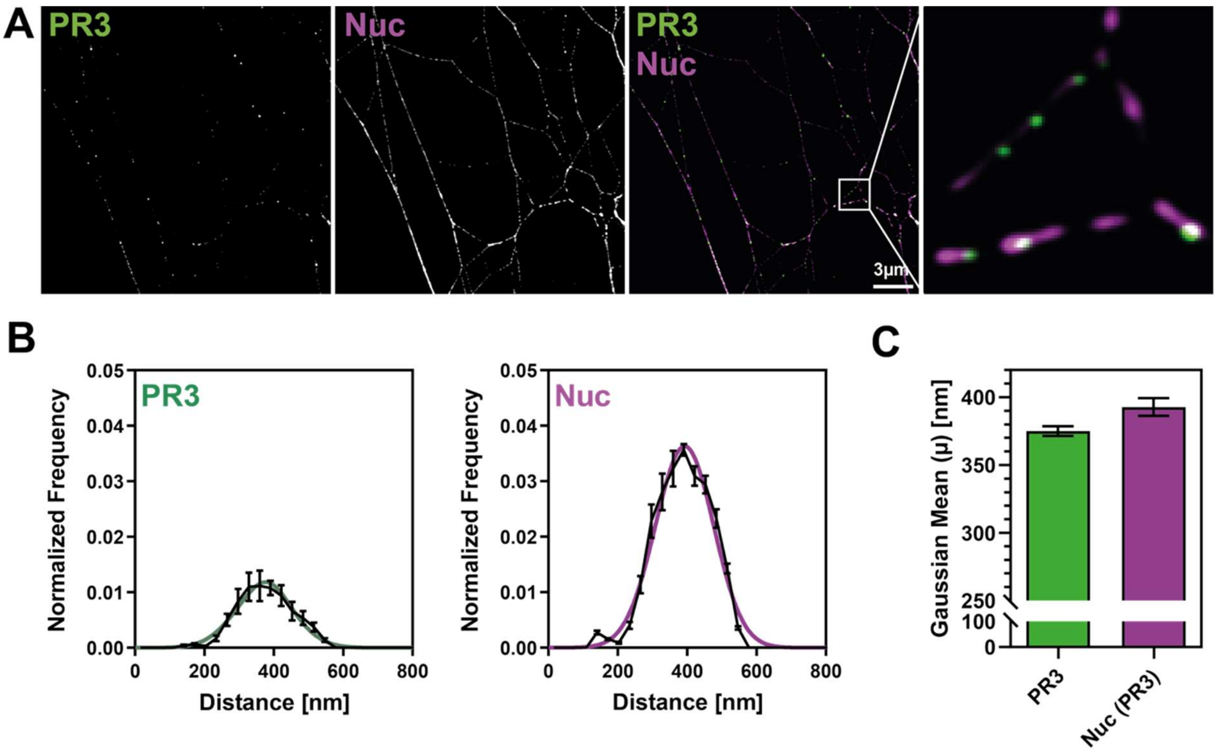


**Supporting Figure S2: SIM of PR3 and nucleosomes on NETs.** Co-labeling of PR3 and NUC on NETs. **(a)** Representative images. Scale bar = 3 µm, box = 2 × 2 µm. **(b)** Periodicity histogram of the proteins displayed in (a) with average of Gaussian fits (colored lines). Data from 5 independent donors; analyzed NET fragments (1.5 µm each): 25,830. Both PR3 and Nuc show strong periodicity, as indicated by the high peak of the periodicity histograms. Means ± SEM (black lines). Centers from Gaussian fits (b) were plotted in **(c)** and reveal similar gaussian distributions for PR3 and Nuc. Means ± SEM.

**Supporting Figure S3**. Dual-color controls (SIM).

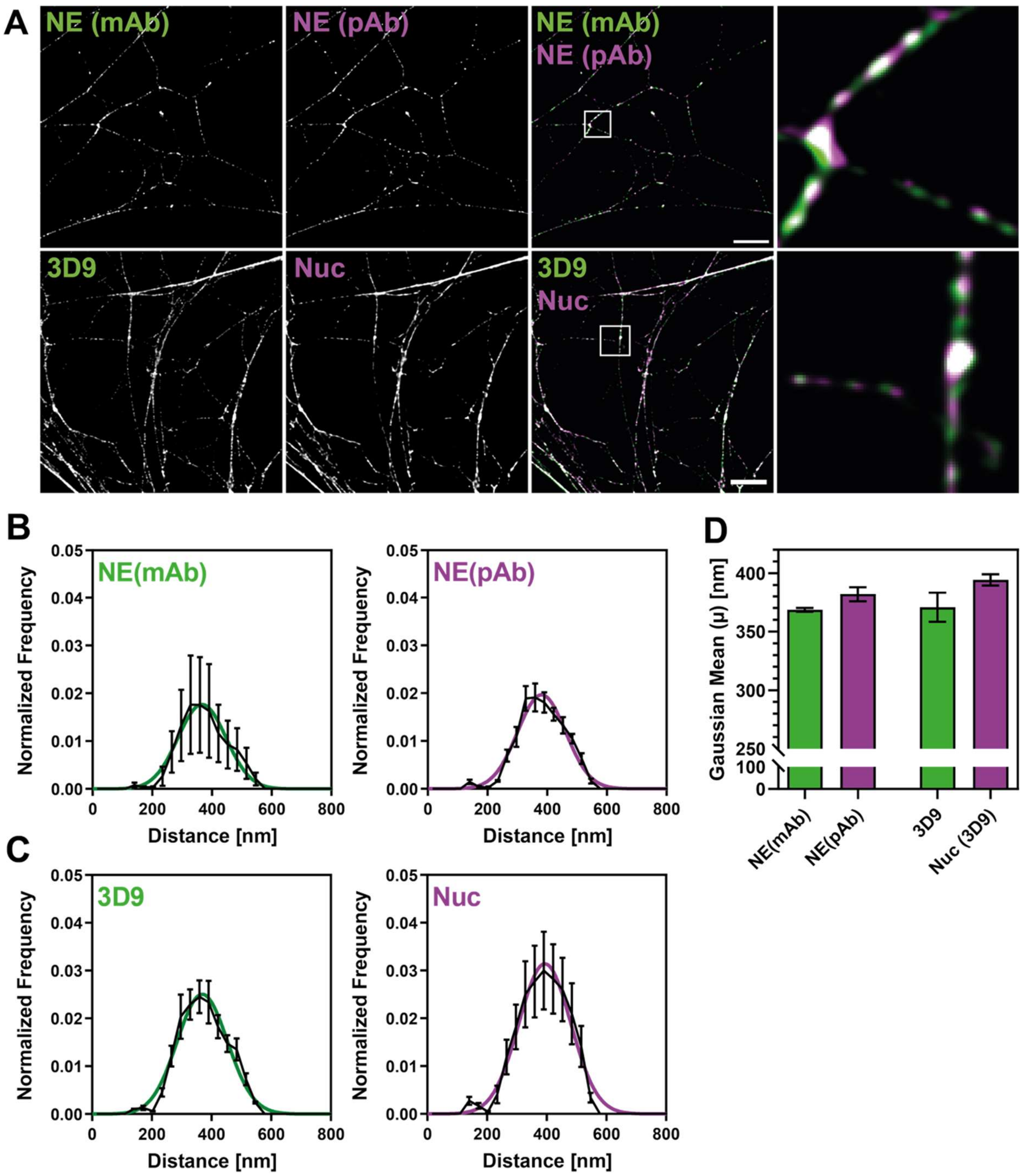


**Supporting Figure S3: Dual-color controls (SIM). (a)** Co-labeling of NE using polyclonal (pAb) and monoclonal (mAb) antibodies (top row), and nucleosomes using the cleaved histone H3 antibody (3D9) and the nucleosome marker (mAB PL2.3) (bottom row). Both NE antibodies co-localize. The same is true for both nucleosomal markers. Analyzed fragments (1.5 µm each): NE (mAb) - NE (pAb) = 14466 from 3 independent donors; 3D9 - Nuc = 54712 from 4 independent donors. Scale bar = 3 µm, boxes = 2 × 2 µm. **(b,c)** Periodicity histogram of the proteins displayed in (a) with average of Gaussian fits (colored lines). Centers from Gaussian fits (b,c) were plotted in **(d)** and served as a measure for the predominant periodicity. Means ± SEM.

**Supporting Figure S4.** SIM of granular and cytosolic proteins.

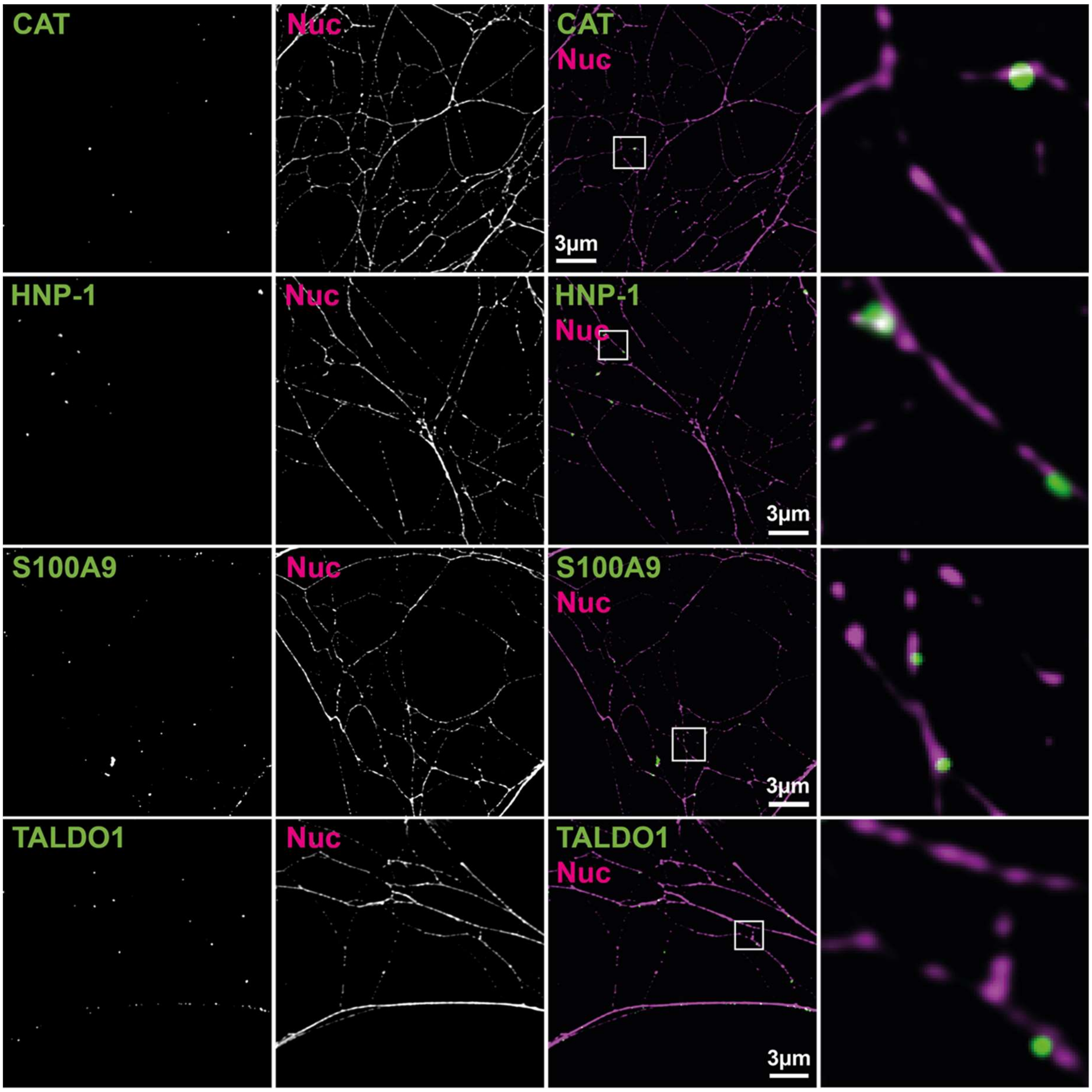


**Supporting Figure S4: SIM of granular and cytosolic proteins.** Co-immunolabeling of granular proteins HNP-1 and CAT, and cytosolic proteins S100A9 and TALDO1 with nucleosomes (Nuc, magenta; mAb PL2.3) on NETs (SIM). All proteins show low-abundant labeling and no obvious periodicity. HNP-1 and S100A9 occasionally form large clusters that co-localize with nucleosomes. Overall colocalization with nucleosomes is low. Scale bar = 3 µm, boxes = 2 × 2 µm.

**Supporting Figure S5.** Periodicity analysis of granular and cytosolic proteins (SIM).

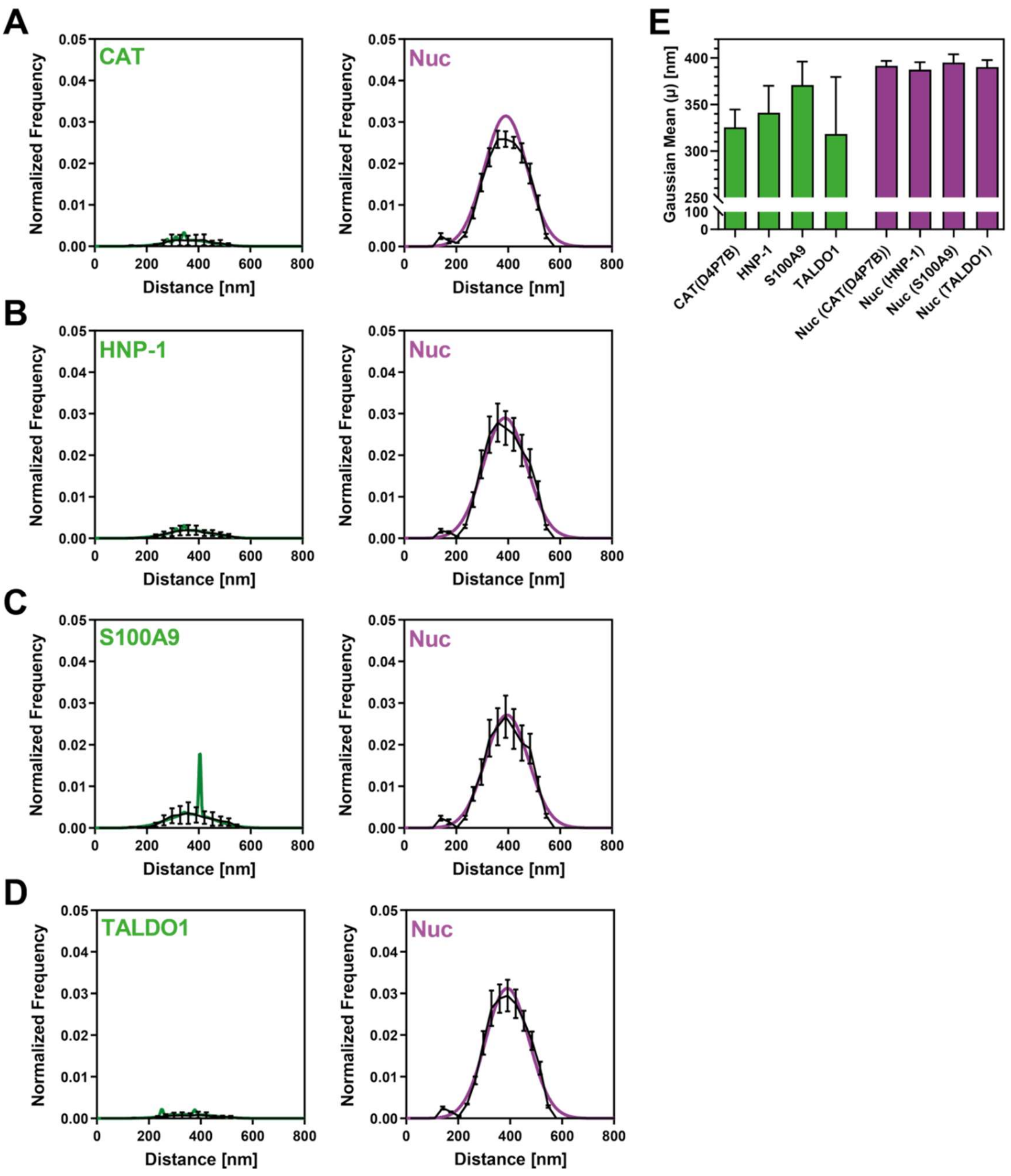


**Supporting Figure S5: Periodicity analysis of granular and cytosolic proteins (SIM). (a-d)** Periodicity histograms of granular (HNP-1, CAT) and cytosolic (S100A9, TALDO1) neutrophil proteins shown in Sup Figure 4, with averages of Gaussian fits (colored lines). All proteins exhibit low periodicity, as indicated by weak peak in the periodicity histogram. Data from four independent donors; analyzed NET fragments (1.5 µm each): HNP-1 = 70,367; S100A9 = 33,890; CAT = 58,019; TALDO1 = 85,005. Means ± SEM (black lines). **(e)** Centers from Gaussian fits (b,c) were plotted in (a-d) and serve as a measure for the predominant periodicity. Means ± SEM.

**Supporting Figure S6.** Colocalization analysis of NET proteins with nucleosomes (SIM).

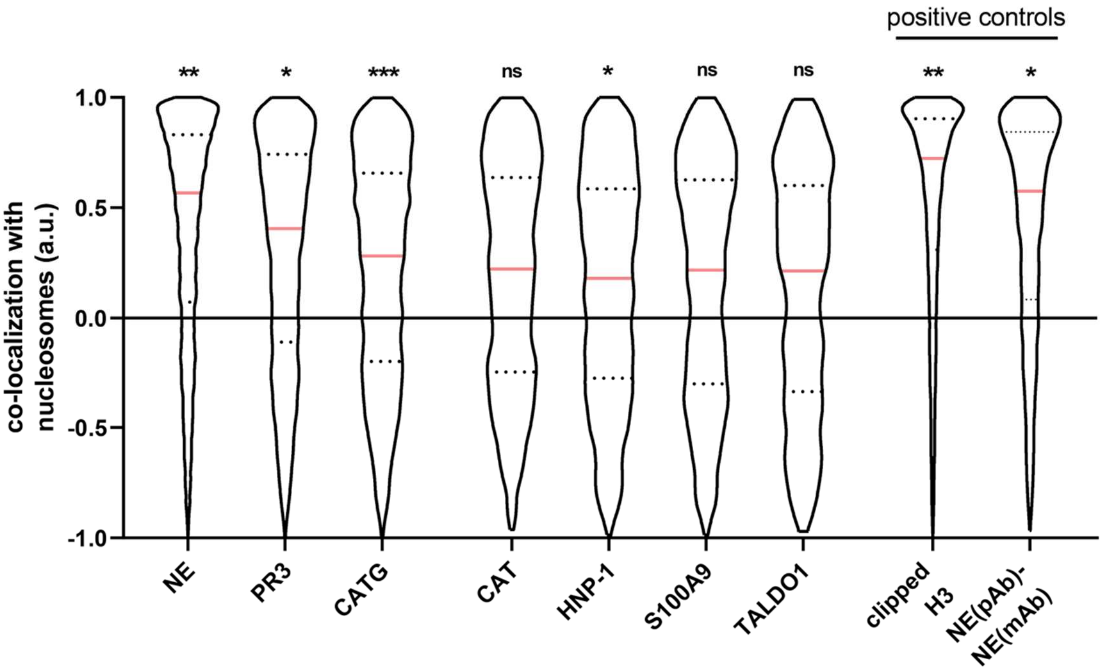


**Supporting Figure S6: Colocalization analysis of NET proteins with nucleosomes (SIM).** NET-associated proteins with nucleosomes and technical positive controls were analyzed. NE, PR3, CATG and HNP-1 and clipped Histone 3 (biological positive control) show significant colocalization with nucleosomes. While NE and PR3 co-localize with nucleosomes to a similar extent as the biological positive control clipped H3 (or pAB NE with mAB NE, technical control), CATG and HNP-1 display only weak colocalization. CAT, S100A9, and TALDO1 exhibit multimodal distributions and no significant colocalization. Median ± quartiles. Data from 3-5 independent donors. Analyzed line profiles and fraction of double-labeled (1.5 µm each): NE = 19,691 (32.6%), PR3 = 6,847 (26.5%), CATG = 7,442 (11.2%), CAT = 2,215 (3.8%), HNP-1 = 1,903 (2.7%), S100A9 = 2,000 (5.9%), TALDO1 = 1,042 (1.2%), clipped H3 = 13,128 (24%), NE (pAb) – NE (mAb) = 2,553 (17.6%). P-values (one-sample t-test and zero, pooled per donor, Bonferroni-corrected): NE: 0.004, PR3: 0.041, CATG: 0.001, CAT: 0.063, HNP-1: 0.004, S100A9: 0.096, TALDO1: 0.081, clipped H3: 0.004, NE (pAb) – NE (mAb): 0.019. * $p < 0.05$, ** $p < 0.01$, *** $p < 0.001$, n.s. not significant.

**Supporting Figure S7:** Multi-color STED microscopy of NETs.

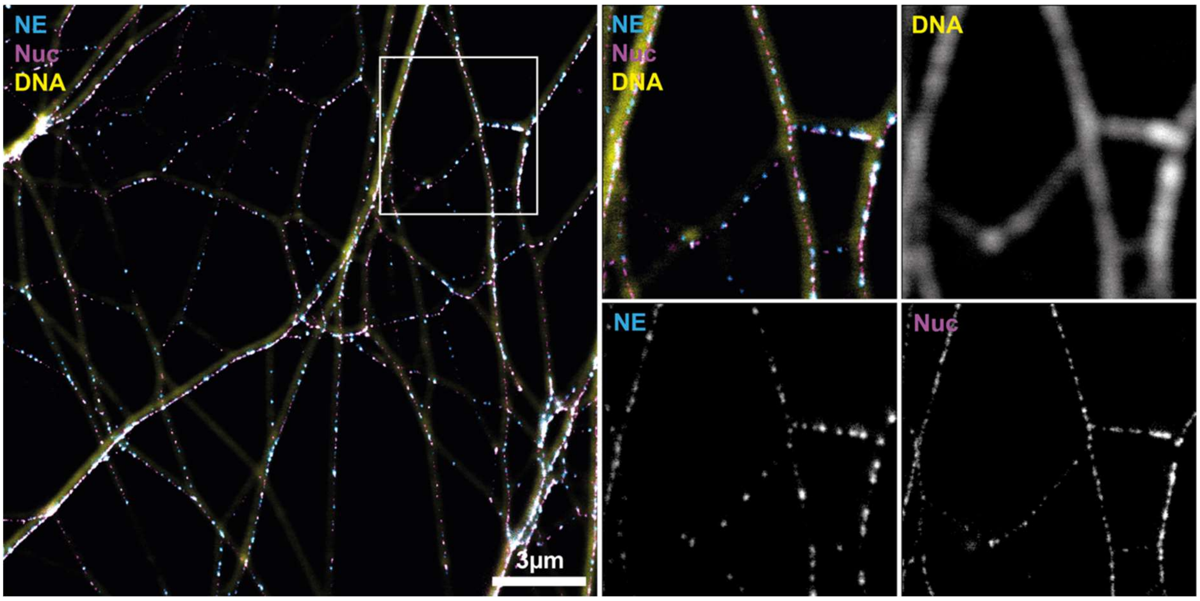


**Supporting Figure S7: Multi-color STED microscopy of NETs on DNA backbone.** The DNA backbone (yellow) was stained with the DNA-intercalating dye YOYO-1 and imaged in confocal mode. NETs were co-labeled with antibodies against NE (cyan) and Nuc (magenta). The overview image (left) shows web-like structure of NETs with individual filaments. NE and Nuc are organized in periodic clusters along the filaments (zoom, right). Diffraction limited confocal images of the DNA channel reveal continues labelling on NET filaments, which was used for NET segmentation. Zoom: 5 × 5 µm.

**Supporting Figure S8.** Multi-color STED microscopy of PR3 and nucleosomes.

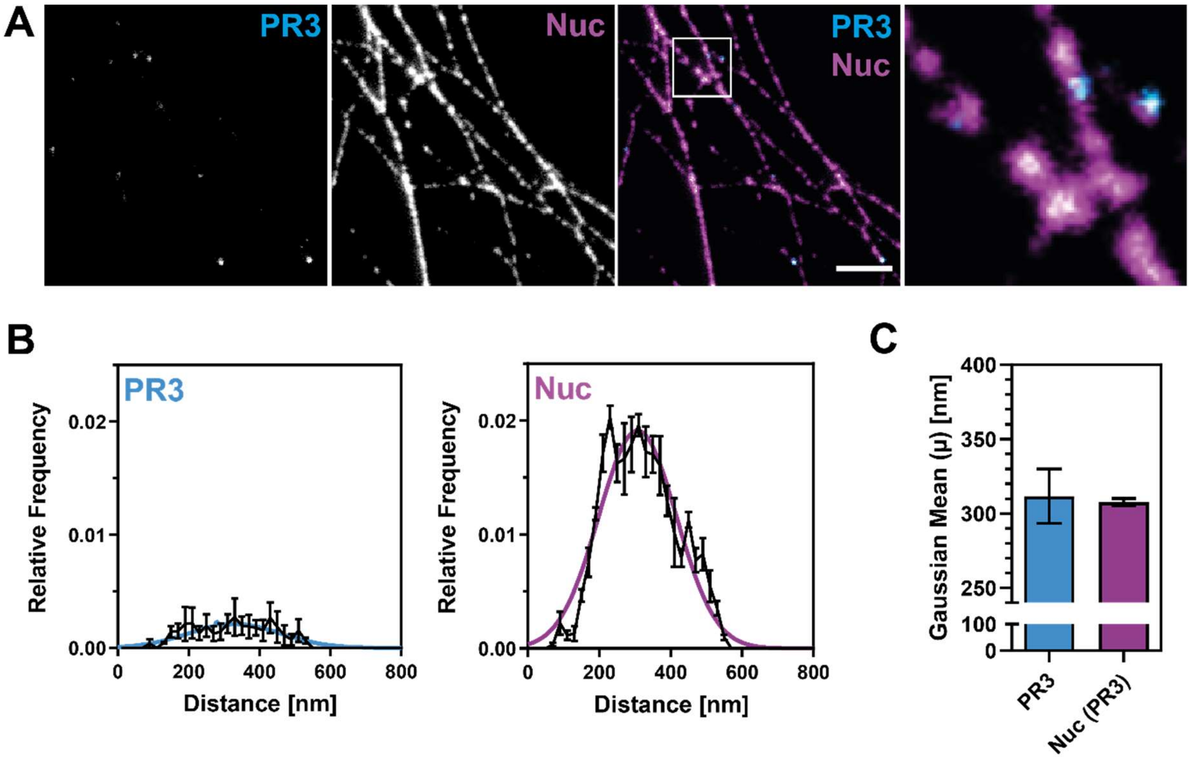


**Supporting Figure S8: Multi-color STED microscopy of PR3 and nucleosomes. (a)** Co-labeling of PR3 and nucleosomes (Nuc) on NETs in STED mode. Scale bar = 2 µm, boxes = 2 × 2 µm. **(b)** Periodicity histogram of the proteins displayed in (a) with average Gaussian fits (colored lines). Both PR3 and Nuc display a prominent peak in the periodicity histogram. Data from three independent donors; analyzed NET fragments: NE = 10,974, CATG = 9,219, PR3 = 7,460. Means ± SEM (black lines). Peaks from the Gaussian fits (b) were plotted in **(c)** and serve as a measure of the predominant periodicity. Means ± SEM.

**Supporting Figure S9.** Colocalization analysis of NET proteins with nucleosomes (STED).

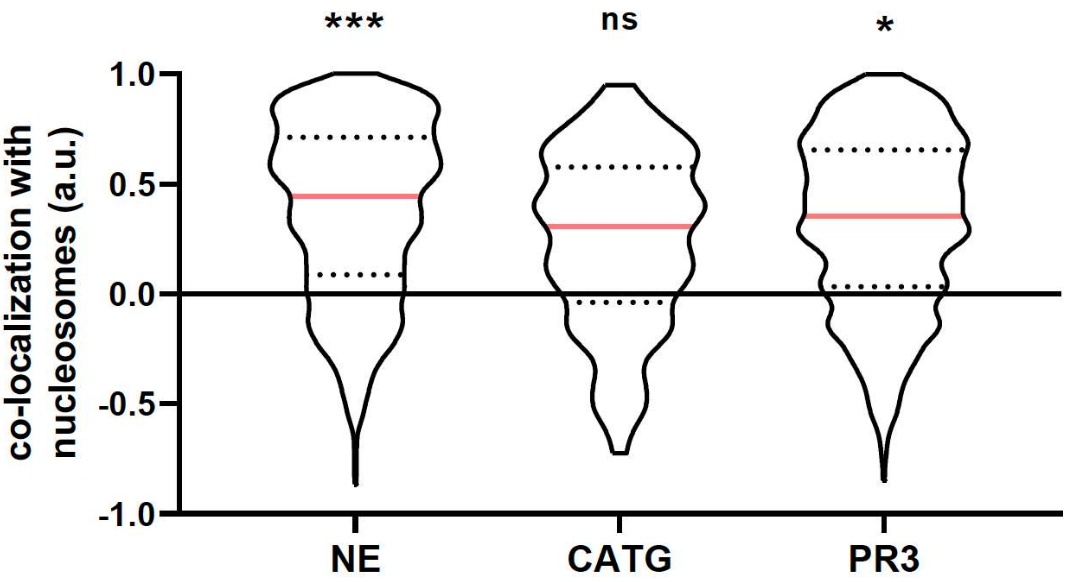


**Supporting Figure S9: Colocalization analysis of NET proteins with nucleosomes (STED).** NE and PR3 show significant colocalization with nucleosomes, while CATG (in contrast to SIM, Figure 6) shows no significant colocalization. Median ± quartiles. Data from three independent donors. Analyzed line profiles and fraction of double-labeled (1.5 µm each): NE = 10,974 (16%), PR3 = 7,460 (7%), CATG = 9,219 (1%). P-values (one-sample t-test and zero, pooled per donor, Bonferroni-corrected): NE: 0.0004%, PR3: 0.043, CATG: 0.056. * $p < 0.05$, *** $p < 0.001$, n.s. not significant.

**Supporting Figure S10.** Schematic binding model of different neutrophil serine proteases.

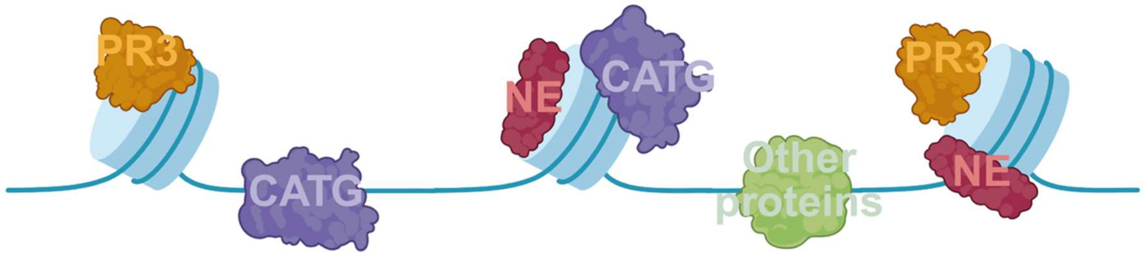


**Supporting Figure S10:** Schematic binding model of different neutrophil serine proteases. The positive colocalization of NE and PR3 with nucleosomes suggests that their binding to NETs is mediated through nucleosome association, in line with the shared periodic distribution. In contrast, CATG displays less pronounced periodicity and reduced colocalization with nucleosomes, indicating a mixed binding mode, potentially involving both nucleosomes and DNA or other components. All other NET-associated proteins tested lacked periodic distribution and except for HNP-1 did not colocalize with nucleosomes, suggesting a nucleosome-independent binding mechanism. Together this suggests that different neutrophil serine proteases use different modes of binding to NETs.

### METHODS

#### Human sample collection and cell lines

Our study was conducted in accordance with the Helsinki Declaration. Anonymous blood donations from the Charité Campus Mitte blood bank as approved by the ethics committee of Charité University Hospital, Berlin, Germany.

#### Neutrophil isolation and cell culture

Blood was collected into EDTA containing tubes, layered 1:1 on Histopaque 1119 (Sigma) followed by centrifugation for 20 min at 800 × g. Plasma and the upper layers of the separated blood, consisting mainly of peripheral blood mononuclear cells, were discarded. The neutrophil-rich pink layer was collected whilst the densest layer consisting of red blood cells was left undisturbed. Neutrophils were washed in PBS containing 0.1% human serum albumin (HSA, Grifols), and further fractionated on a discontinuous Percoll (Pharmacia) gradient consisting of 2mL layers with densities of 1105 g/ml (85%), 1100 g/ml (80%), 1093 g/ml (75%), 1087 g/ml (70%), and 1081 g/ml (65%). Neutrophils were carefully layered on the top of the gradient and centrifuged for 20 min at 800 × g, the interface between the 80% and 85% Percoll layers was collected and washed with PBS containing 0.05% HSA. Neutrophil purity was determined to be > 95% by flow cytometry.

#### NET preparation for super resolution microscopy

Isolated neutrophils were seeded on isopropanol sonicated high-precision coverslips (⌀24mm, 1.5H, Marienfeld, Germany) in 6-well cell culture dishes at a density of $1.5x10^5$ cells/coverslip in 2ml RPMI medium + 0.1% HSA. Cells were allowed to adhere to the coverslip for 15min at 37°C, 5% $CO_2$. NET formation was stimulated by incubation with 100nM PMA (Sigma) for 3:30h at 37°C, 5% $CO_2$. Throughout the NET preparation extra attention was used not to shake or disturb the solution to preserve the delicate structure of NETs. NET formation was validated by light microscopy. Samples were fixed in 3% para-formaldehyde (PFA) (w/v) (Electron Microscopy Sciences) for 12 min, RT by adding 1ml of 3x concentrated PFA to each coverslip within the 6-well cell culture dish. Coverslips were subsequently washed twice with PBS. For immunofluorescent staining, the samples were permeabilized for 10 min with 1% Triton X-100 in PBS at RT and subsequently blocked in fish gelatin/goat serum blocking buffer for 1h, RT. Afterwards, samples were incubated with primary antibodies in fish gelatin/goat serum blocking buffer (0.05% (v/v) Tween 20, 3% (v/v) normal goat serum, 3% (w/v) freshwater fish gelatin, 1% (w/v) BSA in PBS pH 7.5) overnight at 4°C. After two subsequent washes with

PBS, secondary antibodies and DNA dyes were added to coverslips in fish gelatin/goat serum blocking buffer for 1h at RT. Where required, samples were sequentially labeled with primary antibodies from the same host species. Primary antibodies were conjugated to fluorophores by NHS ester labeling according to manufactures protocol (Thermo, # A88068) and added to samples for 1h at room temperature. Samples for STORM were mounted on concave microscopy slides with 100µl oxygen scavenging buffer (0.1 mg/ml GLOX, 0.1 mg/ml HRP, 25mM HEPES, 5% glycerol, 25mM glucose in PBS, pH 6.0) and sealed with a two-component dental imprint material to enable effective photo switching of fluorescent cyanine dyes. Preparation of buffers and mounting of samples was performed immediately before image acquisition. For SIM and STED, coverslips were mounted to HistoBond microscopy slides (Marienfeld, Germany) in three drops ProLonged Glass antifade mountant (ne=1.52, Invitrogen). Mounted NET samples were cured for at least one week at RT before imaging to minimize DNA filament snapping at lower wavelength laser powers.

**SIM microscope setup, image acquisition and reconstruction**

Structured Illumination Microscopy (SIM) was performed using a Zeiss ELYRA 7 Lattice SIM system on an Axio Observer 7 stand. The microscopy was equipped with a Zeiss Plan-Apochromat 63x/1.4 oil DIC M27 objective and HR diode 488nm (500mW), HR DPSS 561nm (500mW) and HR diode 642nm (500mW) laser lines. Simultaneous detection of two channels was achieved by Duolink adaptor, operating two pco.edge 4.2 CL HS sCMOS cameras (6.5µm pixel size, peak QE 82%, liquid cooled) (PCO AG, Kelheim, Germany). Excitation and emission beams were split using the DuoLink SR QUAD filter module with double emission bands and a dichroic mirror (SBS LP 560 with EF BP420-480/BP495-550 & EF570-620 & LP655). Combined with acousto-optic tunable filter (AOTF) switched laser lines this setup enabled quick four-channel image acquisition with the blue and orange as well as the green and red spectra in parallel. Prior to sample imaging, the sCMOS cameras were aligned using the in-built alignment option of the ZEN Black software. Samples were loaded to the microscopy using Immersol 518 F / 30°C (ne=1.518, Zeiss) to match the refractive index of both glass and ProLong Glass (Invitrogen) mountant. Acquisitions were performed with 50-100mW laser powers at 10-30ms exposure for antibody labelling (568nm, 647nm lasers) and 150-200mW laser power at 50ms exposure for YOYO-1 (Invitrogen) DNA labeling (488nm laser), respectively. For image acquisition, 15-20 regions of interest (2048x2048px) were selected per sample and 3.3µm z-stack (30 slices, 0.11µm intervals) were acquired in optimal mode, switching tracks in “frame fast” mode. Structured illumination was set to ‘Lattice SIM’ with a grating of 13 phases. Channel alignment was performed individually for every session using a sample control. NETs were stained for nucleosomes (anti-PL2.3) using secondary antibodies in all channels (Alexa488, Alexa568, STAR635P) and the channel alignment tool (ZEN black software) was used in ‘affine mode’ using markers to generate an alignment matrix. Afterwards, images were reconstructed using the Zeiss SIM2 module in ‘strong fixed’ mode including the channel alignment matrix. Zeiss SIM$^2$ is a proprietary algorithm increasing signal-to-noise ratio, minimizing SIM artifacts.

**STED microscope setup, image acquisition and reconstruction**

Stimulated emission depletion (STED) microscopy was performed on an Abberior STED Facility Line (#FC211501, Abberior GmbH, Göttingen) based on Olympus IX83 Inverted Microscope. The microscope was equipped with a UPXLAPO apochromat 60x/1.42 NA oil objective (Olympus Life Sciences), 405nm/cw (50mW), 485nm/pulsed (~1mW, 40MHz), 561nm/pulsed (~200µW, 40MHz), 640nm/pulsed (~1mW, 40MHz) excitation laser lines and ultra-high power 775nm/pulsed (>2750mW, 40MHz, repetition rate 25-40MHz) STED laser. The laserlines of the microscope were aligned using the Abberior Nanoparticle Autoalignment Slide (NP-3016) and the inbuilt auto-alignment tool in the Inspector software. STED images were acquired in 15x15µm fields-of-view, with a resolution of 20nm/px. NET proteins were imaged using appropriate antibodies and a STAR Orange conjugated secondary antibody by CLSM (30% laser power; 15-22ms) and STED (45% laser power; 30-40ms) with the 561nm laser. Nucleosomes were imaged using the PL2.3 antibody and a STAR 635P conjugated secondary antibody by CLSM (1.5% laser power, 8-15ms) and STED (2.5% laser power, 12-24ms) with the 640nm laser. In STED mode, the 775nm depletion laser was used at 30% laser power and 10ms exposure.

**STORM microscope setup, acquisition reconstruction and line profile generation**

Single molecule localization microscopy (dSTORM) was performed at the Advanced Medical Bio Imaging (AMBIO) facility in the Charité Universitätsmedizin campus Mitte. A Nikon Ti Eclipse based STORM microscope system (N-STORM V3) was used[19]. The EPI-Total Internal Reflection Fluorescence (TIRF) microscope was equipped with an Agilent MLC400 laser box (405nm, 488nm, 561nm, 640nm), a high NA oil objective (100x 1,49N.A.) and a sCMOS camera (Prime 95B, 1024x1024, Photometrics). Optical Filters were from AHF Analysentechnik. The microscope and camera were controlled by NIS Elements software (Nikon) and image analysis was performed in Fiji (ImageJ 1.54c). The dSTORM images were acquired using a highly inclined and laminated optical sheet (HILO) to minimize background staining and facilitated the imaging of thin DNA filaments. Images were acquired at a frame rate of 50 Hz (20 ms) for AlexaFluor 647 labeled samples with a total of 15,000 frames. Single molecules were localized with the Fiji plugin ThunderSTORM [25] using a pixel size of 110nm, 2.5 photoelectrons per A/D count and a base level of 100 photons. Image de-noising was performed using a third order wavelet filter (B-Spline) and molecules were localized using the local maximum method [2*std(Wave.F1)] as a peak intensity threshold. Sub pixel localization was performed using the integrated Gaussian point-spread function (PSF) with a fitting radius of 7 pixel. After the average shifted histogram reconstruction, data was density filtered (5 neighbors, 50nm radius) and were drift corrected. The results were exported as localization lists and reconstructed images (average shifted histogram). Since the single-color dSTORM images lacked DNA labeling required for automated NET detection, filament line profiles were manually selected using ImageJ. The "free-hand selection" tool with a 5-pixel width was employed to choose ROIs representing single NET filaments. These were defined by localizations with a minimal width and by exclusion of filament branches or links. Moreover, to mitigate potential errors arising from data nature or missing localizations, a maximum profile length of 1-2 µm was set. This is in line with previously described autocorrelations for filament analysis[26]. Selected areas were saved as ROIs and line profiles were exported as .csv files.

**Generation of NET filament line profiles**

The z-range encompassing the NET signal was manually selected from the images, followed by the generation of maximum projections id 3d stacks (SIM and STED) from selected areas. Background values were determined from NET-free areas and subtracted from all images per NET preparation before further processing. Images exhibiting low labeling quality in at least one channel were excluded from the analysis, resulting in the removal of approximately 30% of the acquired images. Additionally, non-NET regions, such as remnants of neutrophils, were masked out to ensure that subsequent analyses focused exclusively on the NET structures. While regions in dSTORM images were manually selected, the analysis workflow **"NET-Detection"** from our custom-built **NanoNET toolbox** (https://github.com/ngimber/NanoNET) was employed for automated NET segmentation in SIM and STED images. The NET-detection workflow of NanoNET features a graphical user interface (GUI, Figure 1) to adapt critical parameters and employs ImageJ filters to generate line intensity profiles. Specifically, the DNA backbone channel was first extracted, and intensities were normalized using the "Normalize Local Contrast" function of ImageJ (settings: block radius = 30nm, standard deviation = 90nm, center stretch enabled). Following normalization, Otsu's method for binarization [27] and "Non-local Means De-noising"[28] (settings: sigma = 90nm, smoothing factor = 1) were applied. A morphological closing operation (radius 156 nm) was then performed to refine the segmented structures. The resulting DNA masks were skeletonized, and short skeleton fragments smaller than 420nm) were excluded. NET filaments were subsequently converted into ROIs and exported. Signal intensities were measured along the line profiles in all channels and exported individually for each image. Parameter files are provided in the supplementary materials.

**Correlation analysis**

The auto- and cross-correlation analysis of line profiles was performed using the "Analyze-Profile" workflow from our custom-developed **NanoNET** toolbox (https://github.com/ngimber/NanoNET), running on Jupyter Notebook (Python 3.9.13). The **NET-Analysis** workflow of NanoNET features a graphical user interface (Figure 1) for adjusting key parameters, including pixel size, filament fragment size, number of lags, minimal filament length, and peak prominence threshold, along with options for graphical output and debugging. Specifically, line intensity profiles obtained through the NET-Detection workflow were divided into equally sized fragments (SIM and STED: 1.5 µm, dSTORM 1.0 µm). Auto- and cross-correlation was then computed for each fragment. Measurements were repeated with multiple shifts (lags) introduced for analysis. Correlation values range from -1 (indicating perfect anti-correlation) to 1 (indicating perfect correlation). Shifts corresponding to the predominant periodicity of the sample enhance the correlation, compared to arbitrary shifts. As a result, the predominant periodicities can be identified by locating the first peak in the correlation plot (correlation versus lag; Figure 1). The first peak represents the predominant periodicity of the sample, and following peaks represent integer multiples of the primary periodicity, providing insights into higher-order periodic patterns within the sample. Correlation profiles were smoothed using Gaussian kaernel, with a window size corresponding to the microscope resolution. Peaks were then detected using SciPy find_peaks with a minimum prominence threshold of 0.4. For each protein target, we analyzed the correlation plot of 40,000–100,000 line profile fragments, calculated the first peak, and plotted those in a correlation histogram (bin size corresponds to image pixel size), enabling statistical comparisons between experimental groups. Data outputs of NanoNET included comprehensive correlation result tables, averaged correlation profiles, and lists of extracted periodicities. Parameter files are provided in the supplementary materials. For colocalization analysis, the correlation value at lag = 0 was used and plotted as violin plots (Supporting Figure 7 and 9).

Analyses included data from over four biological replicates, encompassing per protein target. All graphical representations were generated using GraphPad PRISM 5. The complete analysis workflows are available as part of the NanoNET toolbox (https://github.com/ngimber/NanoNET).

### SUPPLEMENTARY TABLES

**Supplementary Table 1.** Primary antibodies

| Name | Antibody | Host | Clonality | Supplier | Catalog # | Conc. / dil. SIM & STED | Conc. / dil. STROM |
|---|---|---|---|---|---|---|---|
| **3D9** | anti-cleaved Histone 3 (3D9) | M | mAb | Brinkmann, Tilley et. al [16] | n.a. | 10µg/ml | |
| **HNP-1** | anti-alpha Defensin 1 | R | pAb | abcam | ab134706 | 1:50 | |
| **CAT (D4P7B)** | Anti-catalase (D4P7B) XP® | R | mAb | Cell Signaling Technology | #12980 | 1:150 | |
| **CATG (EPC)** | Anti-human Cathepsin G | R | pAb | EPC | CA617 | 1:250 | |
| **NE (pAb)** | Anti-Neutrophil Elastase | R | pAb | EMD Millipore | 481001 | 1:100 | 1:100 |
| **NE (mAb)** | Anti-Neutrophil Elastase [NP57] | M | mAb | abcam | ab254178 | 1:50 | |
| **Nuc (PL2.3)** | anti H2A-H2B-DNA (nucleosomes) | M | mAb | Brinkmann, Herlands et al. [29] | n.a. | 2µg/ml | 5µg/ml |
| **PR3** | Anti-Proteinase 3 (anti-serum) | R | pAb | Elastin Products | PR215 | 1:100 | |
| **S100A9** | Anti-S100A9 Antibody | R | pAb | Atlas Antibodies | HPA 004193 | 1:20 | |
| **TALDO1** | Anti-TALDO1 | R | pAb | Atlas Antibodies | HPA 048089 | 1:25 | |

M = mouse; R = rabbit; mAb = monoclonal antibody; pAb = polyclonal antibody; conc. = concentration; dil. = dilution.

**Supplementary Table 2.** Secondary antibodies

| Name | Fluorophore | Supplier | Catalog No | Dilution SIM | Dilution STED | Dilution STORM |
|---|---|---|---|---|---|---|
| anti-ms A568 F(ab')2 | Alexa Fluor 568 | Invitrogen | A11019 | 1:500 | - | - |
| anti-rb A568 | Alexa Fluor 568 | Invitrogen | A-11036 | 1:500 | - | - |
| anti-rb STAR ORANGE | STAR ORANGE | abberior | STORANGE | - | 1:500 | - |
| anti-ms STAR635P | STAR 635P | abberior | ST635P-1001 | 1:500 | 1:500 | - |
| anti-ms A647 F(ab) | Alexa Fluor 647 | Invitrogen | A21237 | 1:500 | - | - |
| anti-rb CF680 F(ab) | CF680 | Sigma | SAB4600362 | - | - | 1:500 |

**Supplementary Table 3.** Dyes & kits

| Name | Type | Catalog # | Supplier |
|---|---|---|---|
| YOYO-1 | Nucleic Acid Dye | ab275546 | abcam |
| CF680 Protein labelling kit | Succinimidyl ester labelling kit | 92220 | Biotium |
| Alexa647 Antibody labelling kit | Succinimidyl ester labelling kit | A88068 | Thermo |
